# neuromaps: structural and functional interpretation of brain maps

**DOI:** 10.1101/2022.01.06.475081

**Authors:** Ross D. Markello, Justine Y. Hansen, Zhen-Qi Liu, Vincent Bazinet, Golia Shafiei, Laura E. Suárez, Nadia Blostein, Jakob Seidlitz, Sylvain Baillet, Theodore D. Satterthwaite, M. Mallar Chakravarty, Armin Raznahan, Bratislav Misic

**Affiliations:** Montréal Neurological Institute, McGill University, Montréal, Canada; Cerebral Imaging Center, Douglas Mental Health University Institute, McGill University, Montréal, Canada; Lifespan Informatics and Neuroimaging Center, Perelman School of Medicine, University of Pennsylvania, PA, USA; Section of Developmental Neurogenomics, National Institute of Mental Health, Bethesda, Maryland, USA

**Keywords:** neuroimaging, annotations, atlas, registration, coordinate system, software, connectomics, gradients

## Abstract

Imaging technologies are increasingly used to generate high-resolution reference maps of brain structure and function. Modern scientific discovery relies on making comparisons between new maps (e.g. task activations, group structural differences) and these reference maps. Although recent data sharing initiatives have increased the accessibility of such brain maps, data are often shared in disparate coordinate systems (or “spaces”), precluding systematic and accurate comparisons among them. Here we introduce the `neuromaps` toolbox, an open-access software package for accessing, transforming, and analyzing structural and functional brain annotations. We implement two registration frameworks to generate high-quality transformations between four standard coordinate systems commonly used in neuroimaging research. The initial release of the toolbox features >40 curated reference maps and biological ontologies of the human brain, including maps of gene expression, neurotransmitter receptors, metabolism, neurophysiological oscillations, developmental and evolutionary expansion, functional hierarchy, individual functional variability, and cognitive specialization. Robust quantitative assessment of map-to-map similarity is enabled via a suite of spatial autocorrelation-preserving null models. By combining open-access data with transparent functionality for standardizing and comparing brain maps, the `neuromaps` software package provides a systematic workflow for comprehensive structural and functional annotation enrichment analysis of the human brain.

## INTRODUCTION

Imaging and recording technologies such as magnetic resonance imaging (MRI), electro- and magnetoencephalography (E/MEG), and positron emission tomography (PET) are used to generate high-resolution maps of the human brain. These maps offer insights into the brain’s structural and functional architecture, including grey matter morphometry [9, 17], myelination [13, 28, 36, 88], gene expression [3, 33], cytoarchitecture [66], metabolism [72], neurotransmitter receptors and transporters [8, 32, 55, 94], laminar differentiation [83], intrinsic dynamics [24, 52, 67] and evolutionary expansion [34, 60, 87, 91]. Such maps are increasingly shared on open-access repositories like NeuroVault [29] or BALSA [77], which, collectively, offer a comprehensive multimodal perspective of the central nervous system. However, these data-sharing platforms are restricted to either surface or volumetric data, and do not integrate standardized analytic workflows.

If researchers generate novel brain maps in their work—such as task fMRI activations or case-control cortical thickness contrasts—how can they interpret them? Ideally there should be a way to systematically compare and contextualize new maps with respect to existing structural and functional annotations, using rigorous statistical methods. In adjacent fields, such as bioinformatics, there already exist multiple widely-used computational methods for functional profiling and pathway enrichment analysis of gene lists [6, 61]. A comparable structural and functional enrichment tool for neuroimaging would have to support three specific capabilities: (1) a method for generating high-quality transformations across multiple coordinate systems, (2) a curated repository of brain maps in their native space, and (3) a method for estimating map-to-map similarity that accounts for spatial autocorrelation.

Creating brain maps frequently requires collating and aggregating data from many individuals, a multi-step procedure involving myriad methodological decisions. One critical step in this process, however, is the transformation of individual data to a common coordinate system [19, 74]. This transformation is often performed to account for anatomical differences between individual subjects prior to group aggregation, and makes derived maps more comparable across datasets. Data collected from MRI are traditionally represented as volumetric images and are therefore commonly transformed to a standard “population” image in volumetric space (e.g., the MNI ICBM 152 template; [22, 49]); however, standardized triangular (i.e., “surface”) meshes are increasingly used to represent data as well (e.g., the fsaverage, fsLR, and CIVET surfaces; [20, 21, 23, 76]). Ultimately, how researchers choose to represent their data can have important impacts on their analyses, and use of both volumetric and surface-based coordinate systems remains prevalent in the literature.

Transforming individual, subject-level data between different representations and coordinate systems is nontrivial and has been the focus of significant research over the past several decades [5, 15, 30, 54, 63, 64, 71, 89, 93]. Comparatively less attention has been given to generating standardized transformations between coordinate systems for group-level or averaged brain maps [42, 43, 57, 90]. However, as researchers continue to produce and share new maps, there is a growing need to both implement robust and accurate group transformations between coordinate systems, and examine whether and how choice of coordinate system may impact comparisons between maps.

In the current report we introduce a new open-access Python toolbox, `neuromaps`, to enable researchers to systematically share, transform, and compare brain maps (Fig. 1). First, we generate a set of group-level transformations between four standard coordinate systems widely used in neuroimaging and integrate them via a set of accessible, uniform interfaces. Next, we curate over 40 reference brain maps from literature published over the past decade to facilitate contextualizing novel brain annotations. Finally, we implement spatial autocorrelation-preserving null models for statistical comparison between brain maps that will help researchers perform standardized, reproducible analyses of brain maps. Collectively, this represents a first step towards creating systematized knowledge and rapid algorithmic decoding of the multimodal multiscale architecture of the brain.

**Figure 1.**
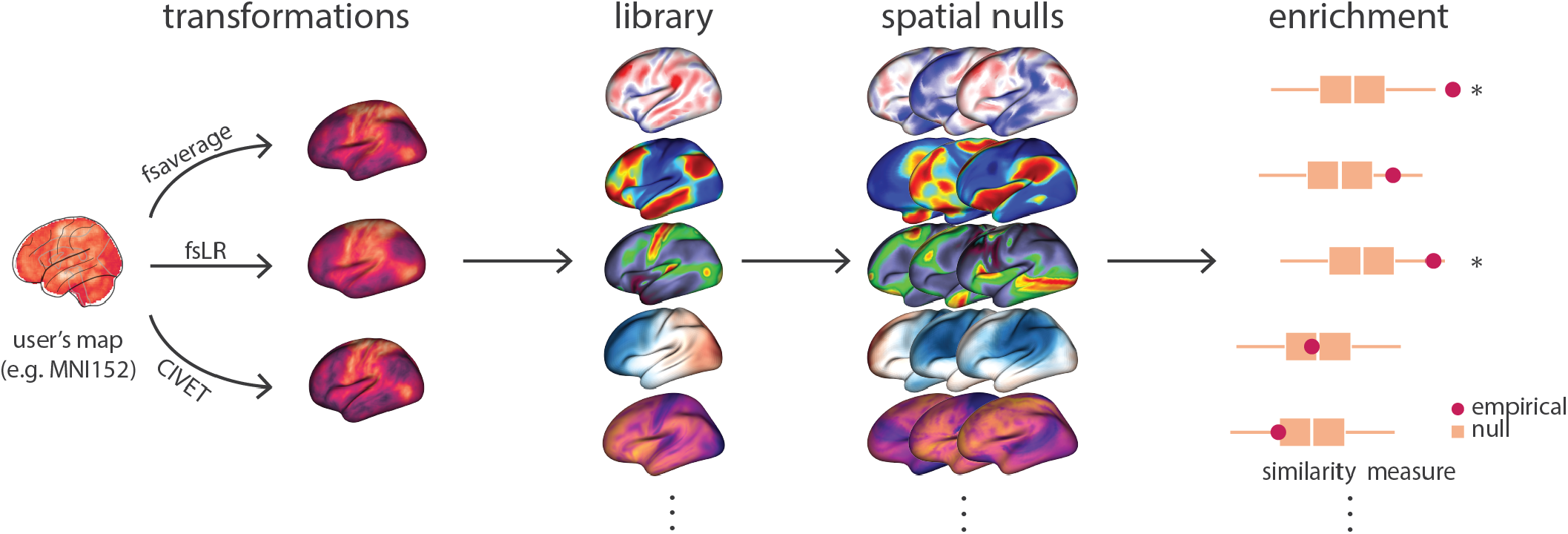
The neuromaps toolbox functionality |. The `neuromaps` software package features a method for generating high-quality transformations across multiple coordinate systems, a curated repository of brain maps in their native systems, and a method for estimating map-to-map similarity that accounts for spatial autocorrelation. For example, user may provide a novel brain map from their own empirical data (e.g. in MNI-152 space), transform it to multiple coordinate systems, and compare it against a library of gradients from the published literature, using spatial autocorrelation preserving null models.

## METHODS AND MATERIALS

### Code and data availability

All code used for data processing, analysis, and figure generation is available on GitHub (https://github.com/netneurolab/markello_neuromaps) and directly relies on the following open-source Python packages: BrainSMASH [14], BrainSpace [82], IPython [59], Jupyter [41], Matplotlib [38], NiBabel [11], Nilearn [1], NumPy [56, 73], Pandas [50], PySurfer [86], Scikit-learn [58], SciPy [81], Seaborn [85], and SurfPlot (https://github.com/danjgale/surfplot). Additional software used in the reported analyses includes CIVET (v2.1.1, http://www.bic.mni.mcgill.ca/ServicesSoftware/CIVET [2]), FreeSurfer (v6.0.0, http://surfer.nmr.mgh.harvard.edu/ [21]), and the Connectome Workbench (v1.5.0, https://www.humanconnectome.org/software/connectome-workbench [45]).

### The neuromaps toolbox

Source code for `neuromaps` is available on GitHub (https://github.com/netneurolab/neuromaps) and is provided under the Creative Commons Attribution-NonCommercial-ShareAlike 4.0 International License (CC-BY-NC-SA; https://creativecommons.org/licenses/by-nc-sa/4.0/). We have integrated `neuromaps` with Zenodo, which generates unique digital object identifiers (DOIs) for each new release of the toolbox. Researchers can install `neuromaps` as a Python package via the PyPi repository (https://pypi.org/project/neuromaps), and can access comprehensive online documentation via GitHub Pages (https://netneurolab.github.io/neuromaps).

### Human Connectome Project

Generating transformations between coordinate systems requires high-quality data from a large cohort of individuals; for the transformations in the `neuromaps` toolbox we use data from the Human Connectome Project (HCP [78]). Raw T1- and T2-weighted structural MRI data were downloaded for *N* = 1113 subjects from the HCP S1200 release. After data processing and omitting subjects that did not successfully complete the CIVET processing pipeline, *N* = 1045 subjects remained.

#### HCP processing pipeline

All structural data were preprocessed using the HCP minimal preprocessing pipelines [27, 78]. Briefly, T1- and T2-weighted MR images were corrected for gradient nonlinearity, and when available, images were co-registered and averaged across repeated scans for each individual. Corrected T1w and T2w images were co-registered and cortical surfaces were extracted using FreeSurfer 5.3.0-HCP [17, 20]. Subject-level surfaces were aligned using a multimodal surface matching (MSMAll) procedure [25].

#### CIVET processing pipeline

Images were separately processed with the `minc-bpipe-library` (https://github.com/CoBrALab/minc-bpipe-library), which performs N4 bias correction, cropping of the neck region, and brain mask generation. Outputs of `minc-bpipe-library` were then processed through the CIVET pipeline (v2.1.0 [2]), which performs non-linear registration to the MNI ICBM 152 volumetric template, cortical surface extraction, and registration of subject surface meshes to the MNI ICBM 152 surface template. Due to CIVET processing failures *N* = 68 subjects were omitted from further analysis.

### Standard coordinate systems

Here we briefly describe the four standard coordinate systems (one volumetric and three surface) considered in the current report. Although other coordinate systems are used in neuroimaging research, these four arguably represent the most commonly-used systems in the published literature.

#### The MNI-152 system

A significant body of work has been dedicated to explaining what is meant when researchers refer to “MNI 152 space”, as several variations of this space exist depending on researchers’ choice of template [10]. In addition to variations on the MNI-152 template, there exist many other MNI spaces which differ from one another enough to impact downstream analyses [44]. Here we use the MNI-152 space as defined by the template from the Minn/Wash-U Human Connectome Project group [78], which is a variation of MNI ICBM 152 non-linear 6th generation symmetric template (identical to the MNI template provided with the FSL distribution [39]). This template was selected because it is the default template in HCP processing pipelines, of which some were used to generate transformations between coordinate systems. This template was created by averaging the T1w MRI images of 152 healthy young adults that had been linearly and non-linearly (over six iterations) transformed to a symmetric model in Talaraich space.

#### The fsaverage system

The fsaverage system, used by FreeSurfer, represents data on the “fsaverage” template, a triangular surface mesh created via the spherical registration of 40 individuals using an energy minimization algorithm to align surface-based features (e.g., convexity; [20, 21]). In current distributions of FreeSurfer there are five scales of the fsaverage template (fsaverage and fsaverage3-6), ranging in density from 642–163,842 vertices per hemisphere. The fsaverage system is roughly aligned to the space of the MNI-305 volumetric system.

#### The fsLR system

Proposed by Van Essen et al. [76], the fsLR coordinate system was created to overcome perceived shortcomings of the fsaverage system: namely, hemispheric asymmetry. That is, the left and right hemispheres of the fsaverage surface are not in geographic correspondence, such that vertex A in the left hemisphere does not correspond to the same brain region as vertex A in the right hemisphere. Researchers used landmark surface-based registration to align the two hemispheres of the fsaverage surface with a common hybrid target. fsLR templates are available in densities ranging from 32,492–163,842 vertices per hemisphere. The fsLR system is roughly aligned to the space of the MNI-152 volumetric system.

#### The CIVET system

The coordinate system used by the CIVET software is a surface reconstruction of the volumetric MNI ICBM 152 non-linear 6th generation template [23]. In its most commonly-used format each hemisphere is represented with 41,962 vertices; a high-resolution version with 163,842 vertices per hemisphere is also available. Because this system is derived from the volumetric MNI template, it ensures that aligned surfaces have good correspondence with volumetric images in the MNI-152 system.

### Generating transformations between systems

Although there are numerous methods for transforming data between coordinate systems, high-quality mappings for group-averaged data are limited [42, 43, 90]. In creating the `neuromaps` toolbox, we used two previously-validated frameworks to generate transformations between all four standard coordinate systems described above (Fig. 2).

**Figure 2.**
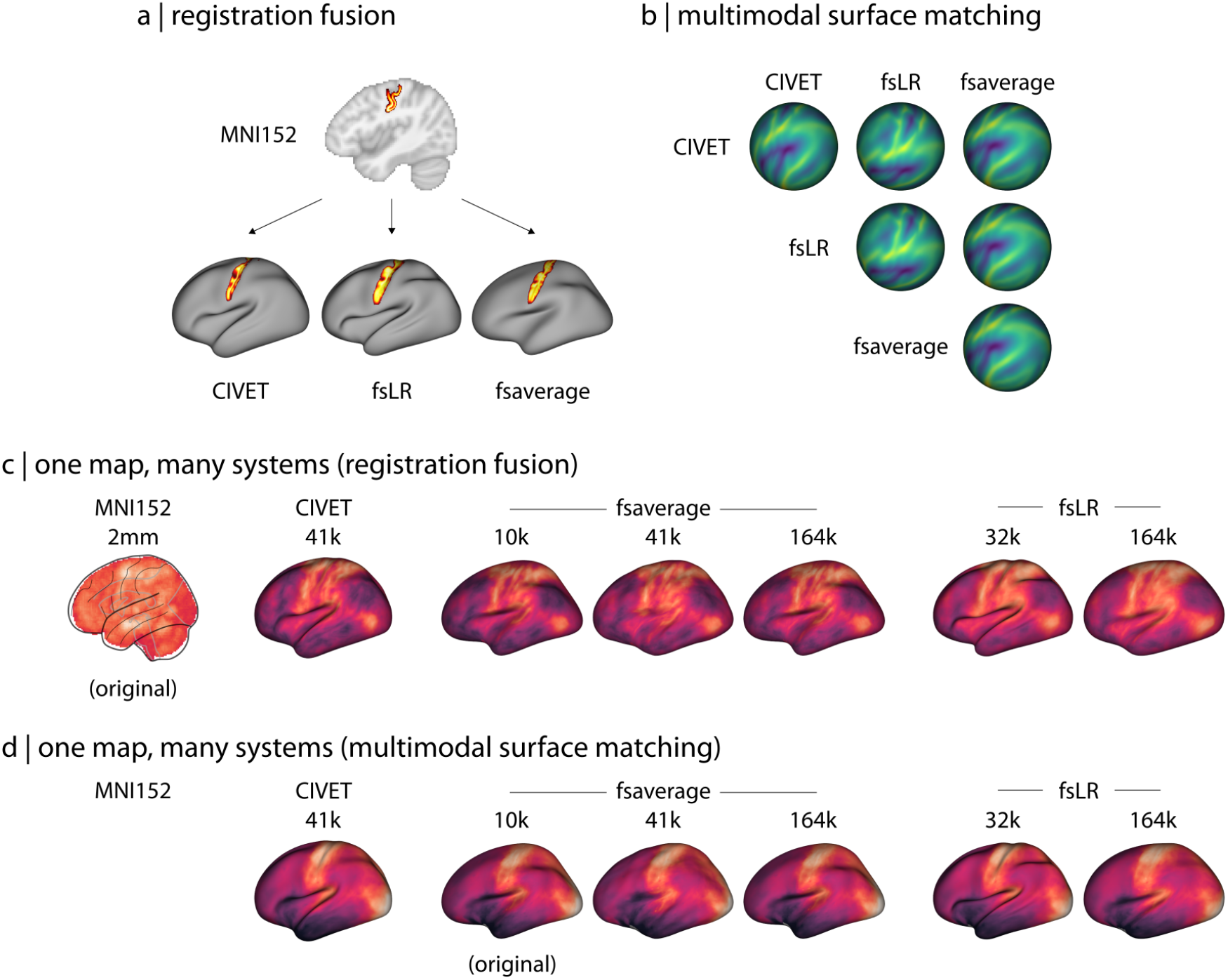
Transformations between coordinate systems |. (a) Registration fusion provides a framework for directly projecting group-level volumetric data to the surface. Here, a probabilistic atlas for the central sulcus [90] has been depicted independently on the CIVET (41k), fsLR (32k), and fsaverage (164k) surfaces. (b) Multimodal surface matching (MSM) provides a framework for aligning spherical surface meshes. Here, we show sulcal depth information (originally defined in fsLR space) on spherical meshes that are aligned across the different coordinate systems, where each row represents a different coordinate system and each column represents the space to which that system is aligned. (c) Example of a volumetric brain map (the first principal component of cognitive terms from NeuroSynth [92]) that has been transformed to all surface-based coordinate systems using alignments derived from registration fusion. (d) Example of a surface brain map (the first principal component of gene expression from the Allen Human Brain Atlas [33]) that has been transformed to all other surface-based coordinate systems using alignments derived from MSM. Note that because the original data are represented on the cortical surface, transformation to volumetric space is ill-defined and therefore not shown here.

#### Registration fusion framework

Originally proposed by Buckner et al. [12] and later developed by Wu et al. [90], registration fusion is a framework for projecting data between volumetric and surface coordinate systems. In its most well-known implementation, researchers used data from the Brain Genomics Superstruct Project (GSP [35]) to generate nonlinear mappings between MNI152 space and the 164k fsaverage surface [90].

Registration fusion works by generating two sets of mappings for a group of subjects: (1) a mapping between each subject’s native image and MNI152 space, and (2) a mapping between each subjects native image and fsaverage space. These mappings are concatenated (MNI152 to native to fsaverage) and then averaged across subjects, yielding a single, high-fidelity mapping that can be applied to new datasets.

Here we generated mappings via registration fusion between the MNI152 volumetric and the fsaverage, fsLR, and CIVET surface-based coordinate systems using data from the HCP (*Methods: Data*). All mappings used functionality from the Connectome Workbench [45] rather than FreeSurfer to ensure standardization of methodology irrespective of target coordinate systems.

##### MNI152 to CIVET

Unlike for fsaverage and fsLR surfaces, CIVET surfaces are extracted from subject T1w volumes after the images have been transformed to the standard MNI152 system. As such, there is no need to generate composite mappings for CIVET surfaces as for the other coordinate systems. Instead, we simply computed the mapping from each subject’s MNI152-transformed T1w volume to the subject’s native CIVET surface, and then applied the CIVET-generated surface resampling to register the mapping to the CIVET standard template system. These mappings were then averaged across subjects to generate a single, group-level transformation.

##### fsaverage/fsLR/CIVET to MNI152

Although every surface vertex has a corresponding voxel representation in volumetric space, not every voxel has a corresponding vertex representation in surface space. As such, generating transformations from the surface coordinate systems to the MNI152 volumetric system cannot yield a “dense” output map. When Wu et al. [90] proposed the current registration fusion framework they adopted a nearest-neighbors, ribbon-filling approach to handle this shortcoming; however, this is only a viable approach when applied to label data (i.e., integer-based parcellation images). We reproduce their approach for completeness, but caution against the application of surface-to-volume projections for continuous data and omit such projections from our analyses.

#### Multimodal surface matching (MSM) framework

The multimodal surface matching (MSM) framework, initially developed by Robinson et al. [63] and later expanded in Robinson et al. [62], aims to align surfaces defined on different meshes using information from various descriptors of brain structure and function. This procedure has been previously used to generate mappings between the fsaverage and fsLR coordinate systems.

Here, we used MSM to generate a mapping between the CIVET and fsLR systems by aligning HCP subject data processed through the CIVET pipeline with the same data processed through the HCP processing pipeline. As MSM requires input data be provided on spherical surface meshes—a representation not produced in the standard CIVET pipeline—we used FreeSurfer functionality to generate spherical mesh representations and sulcal depth information for each subject’s CIVET-derived white matter surfaces. We used these spherical meshes and sulcal depth measurements to drive alignment between the CIVET and fsLR systems via two rounds of the MSM procedure. The first round was used to generate a rotational affine transform to align gross features of the CIVET and fsLR systems; the generated affines were averaged across all subjects and used to seed a second round of finer-resolution alignment, similar to the procedure described in Robinson et al. [62]. The final, aligned subject-level spherical surfaces defined in the CIVET system were averaged to create a single, group-level surface that could be used in future transformations.

The CIVET-to-fsaverage mapping was generated as the composite of the transformations between the CIVET- and-fsLR and fsLR-and-fsaverage systems.

#### Parcellations

Performing analyses at the voxel or vertex level can be computationally intensive. The `neuromaps` software package can be extended to parcellated data and also integrates easy-to-use tools for parcellating volumetric and surfacic data. The base parcellating function assumes that the given parcellation indexes each region with a unique value, where values of 0 are ignored. However, helper functions are provided to flexibly handle alternative parcellation formats, for example, where both hemispheres are indexed with the same range of values.

### Published brain maps

We curated a selection of brain maps from the published literature of the past decade (Fig. 3). Maps were obtained in their original coordinate system. A complete list of maps and their coordinate systems is provided in Table S2. The `neuromaps` data repository will continue to evolve and supports a single-function data contribution pipeline for the neuroimaging community to share novel brain annotations in volumetric or surface-based coordinate spaces.

**Figure 3.**
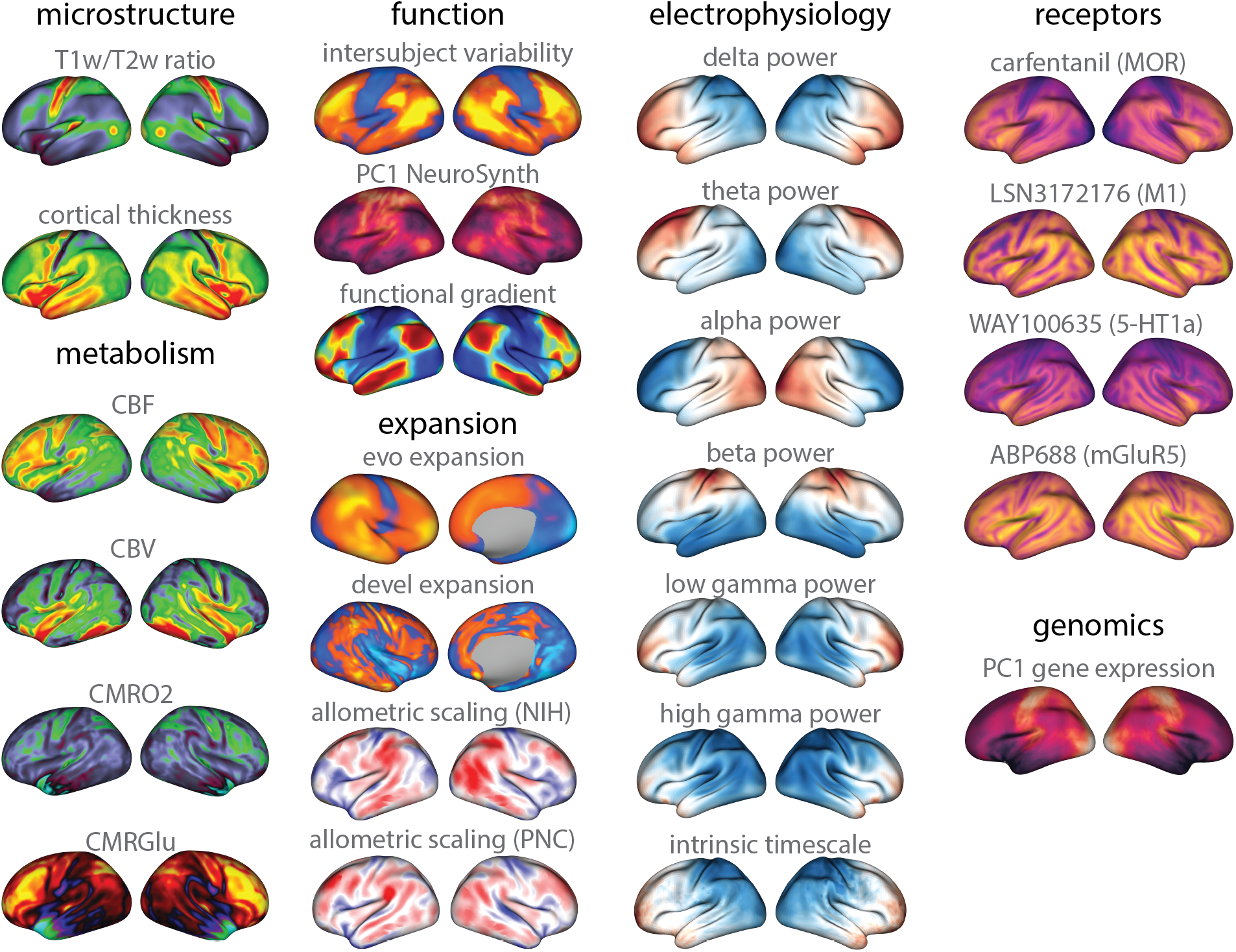
Brain maps from the published literature |. Collection of brain maps obtained from the published literature over the past decade that are currently available in the `neuromaps` distribution. The maps capture the normative multiscale structural and functional organization of the brain, including molecular, cellular, metabolic, and neurophysiological features. Refer to Table S2 for more information on the coordinate system, resolution, and original publication for each brain map. Colormaps were chosen to maximize similarity with how data were represented in the original publication. Note that two of the maps (second column: evo and devel expansion) only have data for the right hemisphere, and the MEG timescale is log-transformed. Additionally, only a selection of four of the 36 neurotransmitter receptor maps are shown here [32, 40, 53, 65, 68]. The repository is continuously updated and the most current list of maps can be found at: https://netneurolab.github.io/neuromaps/. The `neuromaps` toolbox supports a data contribution pipeline and will continue to expand.

A portion of these maps were originally defined in coordinate systems that are now deprecated. We briefly describe the transformations we used to project these maps to one of the current standard coordinate systems (see *Methods: Standard coordinate systems*).

#### PALS-TA24 to fsLR

Data obtained from Hill et al. [34] were originally aligned to a study-specific PALS-TA24 template (derived using a similar landmark-based procedure to the PALS-B12 template; [75]), which has been supplanted by the fsLR coordinate system (see *Methods: The fsLR system*). In order to project data from the PALS-TA24 template to the fsLR system we applied the deformation map provided by the original researchers for transforming data between these spaces using nearest neighbors interpolation.

#### CIVET v1 to v2

The maps obtained from Reardon et al. [60] were originally created using surface templates from CIVET v1.1.12; however, with the release of CIVET v2.0.0 in 2014 the population surface templates provided with the CIVET distribution were updated, effectively deprecating the older templates. In order to project data from the CIVET v1.1.12 templates to the CIVET v2.0.0 templates we used a nearest neighbors interpolation, matching vertex coordinates in the newer template to coordinates in the older template and assigning the value corresponding to the closest vertex [60].

### Spatial null frameworks

Recent research has consistently highlighted the importance of spatially-constrained null models when statistically comparing brain maps [4, 14, 48]. The `neuromaps` software package integrates nine different spatial null frameworks, described in Markello and Misic [48]. These include six spatial permutation models and three parametrized data models which, collectively, can be constructed for surface-based, volumetric, and parcellated data [4, 7, 13, 14, 16, 79, 80, 82]. Note that four of the null models are adaptations of the original spatial permutation framework proposed by Alexander-Bloch et al. [4] when applied to parcellated data [7, 16, 79, 80]. These frameworks differ in how they reassign the medial wall—for which most brain maps contain no data—whether that be by discarding missing data [7, 16], ignoring the medial wall entirely [79], or reassigning missing data to the nearest parcel [80]. The three parametrized data models circumvent spatial rotations by applying generative frameworks such as a spatial lag model [13], spectral randomization [82], or variogram matching [14].

For analyses in the current report using surface-based coordinate systems we apply the procedure proposed by Alexander-Bloch et al. [4]; for analyses using volumetric systems we apply the procedure described by Burt et al. [14]. Null distributions were systematically derived from 1,000 null maps generated by each framework. The mechanism for each null framework used for analyses in the present work is described briefly below.

#### Spatial permutation null model

The procedure proposed by Alexander-Bloch et al. [4] generates spatially-constrained null distributions by applying random rotations to spherical projections of a cortical surface. A rotation matrix (**R**) is applied to the three-dimensional coordinates of the cortex (**V**) to generate a set of rotated coordinates (**V_rot_** = **VR**). The permutation is constructed by replacing the original values at each coordinate with those of the closest rotated coordinate. Rotations are generated independently for one hemisphere and then mirrored across the anterior-posterior axis for the other.

#### Variogram estimation null model

The procedure described by Burt et al. [14] operates in two main steps: first the values in a given image are randomly permuted, then the permuted values are smoothed and re-scaled to reintroduce spatial autocorrelation characteristic of the original, non-permuted data. Reintroduction of spatial autocorrelation onto the permuted data is achieved via the transformation **y** = |*β*|^1/2^**x**′ + |*α*|^1/2^**z**, where **x**′ is the permuted data, 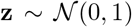 is a vector of random Gaussian noise, and *α* and *β* are estimated via a least-squares optimization between variograms of the original and permuted data.

### Assessing the impact of coordinate system

When transforming two datasets (i.e., a “source” and “target” dataset) defined in distinct coordinate spaces to a common system there are at least three options available: (1) transform the source dataset to the system of the target, (2) transform the target dataset to the system of source, or (3) transform both source and target datasets to an alternate system. If comparisons are being made across several pairs of datasets a fourth option becomes available: (4) always transform the higher resolution dataset to the system of the lower resolution dataset.

To examine whether the choice of coordinate system impacts statistical relationships estimated between brain maps we performed several analyses. First, we transformed a selection of twenty brain maps (see *Methods: Published brain maps*) into every other coordinate system (e.g., fsaverage → fsLR/CIVET/MNI152, fsLR → fsaverage/CIVET/MNI152, and so on). We then correlated every pair of these brain maps according to each of the four possible resampling options described above. When transforming both source and target datasets to an alternate system (option 3, above), we comprehensively tested every target coordinate system and data resolution. Spatial null models were used to assess the significance of all correlations.

## RESULTS

In this report we introduce the `neuromaps` toolbox, an open-access software package designed to streamline and standardize analyses of neuroimaging-derived brain maps (available at https://github.com/netneurolab/neuromaps). The `neuromaps` toolbox provides a uniform interface for transforming brain maps between coordinate systems, contextualizing novel brain maps with respect to canonical structural and functional annotations, and assessing relationships between maps using spatial null models. In the following section we highlight features available in `neuromaps`, demonstrate typical workflows enabled by its functionality, and use `neuromaps` to examine how choice of coordinate system can impact statistical analyses of brain maps.

### The neuromaps data repository

The `neuromaps` toolbox provides programmatic access to templates for four standard coordinate systems: fsaverage, fsLR, CIVET, and MNI-152 (see *Methods: Standard coordinate systems*). For surface-based coordinate systems, we distribute template geometry files, sulcal depth maps, and average vertex area shape files (computed from HCP participants) in standard GIFTI format. For volumetric coordinate systems, we distribute T1-, T2-, and PD-weighted template files, a brain mask, and probabilistic segmentations of gray matter, white matter, and cerebrospinal fluid in standard gzipped NIFTI format.

Beyond template files, the `neuromaps` toolbox offers access to a repository of brain maps obtained from the published literature (Fig. 3; Table S2). These maps were generated using multiple imaging techniques, including magnetic resonance imaging, magnetoencephalography, positron emission tomography and microarray gene expression. Brain maps are provided in the original coordinate system in which they were defined to avoid errors caused by successive interpolation. Collectively, these maps represent more than a decade of human brain mapping research and encompass phenotypes including: the first principal component of gene expression [33], 36 neurotransmitter receptor PET tracer images [32], glucose and oxygen metabolism [72], cerebral blood flow and volume [72], cortical thickness [78], T1w/T2w ratio [25], 6 canonical MEG frequency bands [70, 78], intrinsic timescale [70, 78], evolutionary expansion [34], 3 maps of developmental expansion [34, 60], the first 10 gradients of functional connectivity [46], intersubject variability [51], and the first principal component of Neurosynth-derived cognitive activation [92]. This repository of data is organized by tags and can be easily downloaded directly from `neuromaps`.

The `neuromaps` toolbox makes it easy to contextualize novel brain maps to a range of molecular, structural, temporal, and functional features. This will facilitate an expansion of research questions, allowing researchers to bridge brain topographies across several spatial scales and across disciplines outside of their immediate scope. Additionally, annotations will be regularly added to the repository, resulting in an increasingly rich toolbox. The `neuromaps` toolbox also integrates an easy-to-use workflow for researchers who wish to share new brain maps in any of the four supported coordinate systems; information on contributing new brain maps can be found in the online documentation for the software (https://netneurolab.github.io/neuromaps/).

### Transformations between coordinate systems

Despite the multiscale, multimodal collection of brain phenotypes in `neuromaps`, data cannot be readily compared to one another because they exist in different native coordinate systems. Indeed, a common challenge when relating novel neuroimaging data to the broader literature is finding a common coordinate space or parcellation in which to conduct the analyses. The `neuromaps` module provides transformations between four supported coordinates systems as well as a standardized set of functions for their application. Transformation between volumetric- and surface-based coordinate systems were derived with a registration fusion framework (Fig. 2a; [90]), whereas transformations between surface-based coordinate systems were derived using a multimodal surface matching (MSM) framework (Fig. 2b; [62, 63]). We leverage tools from the Connectome Workbench to provide functionality for applying transformations between surface systems; however, users do not need to interact directly with these Workbench commands. In addition to transforming individual annotations, the `neuromaps` software package includes functionalities that resample images to each others’ spaces. By default, data is only ever downsampled, which ensures `neuromaps` does not artificially create new data. Collectively, the `neuromaps` toolbox facilitates robust transformations between coordinate systems, facilitating the standardization of neuroimaging workflows.

### Assessing relationships between brain maps

The primary goal for transforming maps to a common coordinate system is to statistically compare their spatial topographies. The `neuromaps` software package uses a flexible framework for examining relationships between brain maps, offering researchers the ability to provide their own image similarity metric or function, as well as easy handling of missing data. By default, the primary map comparison workflows use the standard Pearson correlation to test the association between provided maps. The `neuromaps` comparison workflow also integrates multiple methods of performing spatial permutations for significance testing.

Multiple spatial null model frameworks allow for statistical comparison between brain maps while accounting for spatial autocorrelation [4, 7, 13, 14, 16, 79, 80, 82]; however, the implementation of these models varies and, to-date, there has been limited effort to provide a standardized interface for their use. We have incorporated nine null models into the `neuromaps` toolbox, offering a common user interface for each model that can be easily integrated with other aspects of the toolbox. Given the computational overhead of these models, our implementations offer mechanisms for caching intermediate results to enable faster re-use across multiple analyses. Spatial null models are enabled by default in the primary map comparison workflows to encourage their broader adoption. Based on prior work benchmarking the accuracy and computational efficiency of these models [48] we set the non-parametric method originally described in Alexander-Bloch et al. [4] as the default for use with surface data, and the parameterized generative method described in Burt et al. [14] as the default for use with volumetric data.

To demonstrate the utility of `neuromaps`, we analyzed a sample of 20 brain maps from the published literature over the past decade (2011–2021), including two microstructural, four metabolic, three functional, four expansion, six band-specific electrophysiological signal power, and one genomic annotation. We then used `neuromaps` to transform these maps from their original representation to the space defined by each of four standard coordinate systems, for a total of seven different representations (Fig. 3). Finally, we computed the pairwise correlations between all maps in each of the systems and assessed the statistical significance of these relationships using spatial null models (see *Methods: Spatial null frameworks*). The goal of this analysis was twofold. First, we sought to assess the extent to which coordinate transforms could influence map-to-map comparisons. Second, given the growing interest in how these system-level maps or “gradients” are related to one another, we sought to assess patterns of relationships among them [37, 69].

Fig. 4a shows that for most map-to-map comparisons, choice of coordinate system has a minimal effect: correlations between maps on average only change by |*r*| = 0.018. There are also instances in which associations between maps are statistically significant in one coordinate system and not significant in another (Fig. 4c); however, in most cases the p-value for these relationships often fell very close to the statistical alpha (i.e., *p* ≤ 0.05) such that the actual effect size only changed by *r* ≤ 0.10. Across all examined systems, we find that the brain maps tend to form two distinct clusters (Fig. 4b), largely recapitulating previously-established relationships observing anterior-posterior and unimodal-transmodal axes of variation [31, 46, 67]. These results are encouraging, suggesting that transforming brain annotations between different systems generally preserves their relationships.

**Figure 4.**
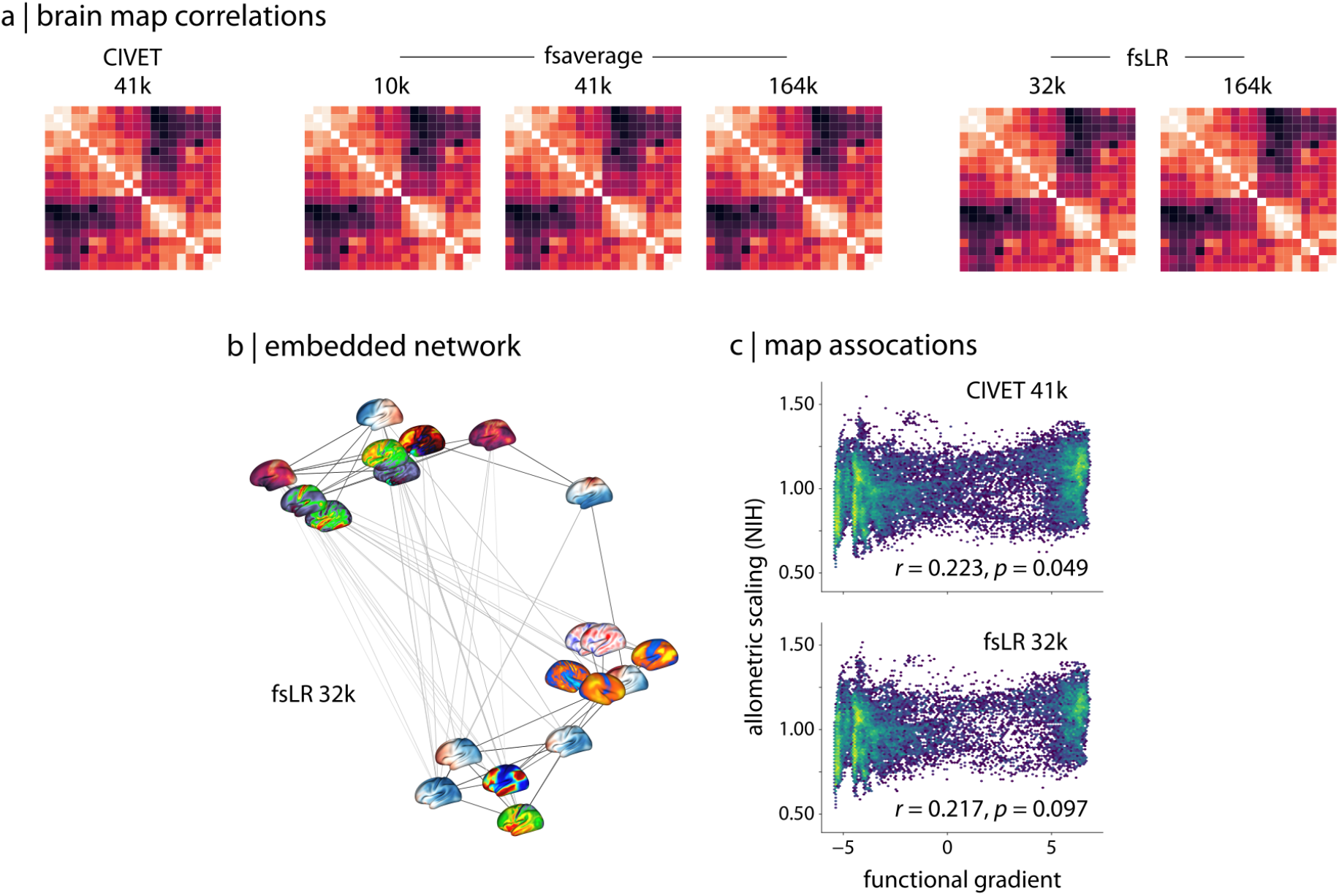
Correlations between brain maps across systems |. (a) Correlation matrices between a selection of twenty brain maps in the `neuromaps` toolbox for each of the surface-based coordinate systems. Because transformations from surface-based to volumetric systems are ill-defined for continuous data we omit those associations. (b) A spring-embedded representation of the correlation matrix among the twenty brain maps, shown here for the fsLR 32k system. (c) An example of two brain maps—the principal functional gradient from Margulies et al. [46] and allometric scaling from Reardon et al. [60]—whose association is significant in one system (CIVET 41k; *r* = 0.223, *p* = 0.049) and not in another (fsLR 32k; *r* = 0.217, *p* = 0.097).

## DISCUSSION

Neuroimaging researchers are constantly generating new datasets that offer unique perspectives into the structural and functional architecture of the human brain. Contextualizing new datasets with multiscale, multimodal brain organization requires both a curated repository of annotations and transparent functionality for transforming and comparing brain maps. Here we introduced the `neuromaps` toolbox which offers both a repository of widely-used brain map datasets as well as a set of standardized methods for transforming them between different coordinate systems and contextualizing novel findings within the broader literature.

Given the proliferation of such datasets in recent years, a large body of work has arisen focused on investigating similarities across brain maps [7, 18, 24, 31, 67, 80, 84]. Indeed, researchers have observed significant concordance in the spatial topology of brain maps derived from a wide variety of phenotypes, suggesting that these maps may reflect a fundamental organizational principle of the human brain. We reproduce this phenomenon in our example analyses, wherein twenty datasets provided with the `neuromaps` toolbox seem to group into two distinct clusters (Fig. 4). One cluster contains maps including the T1w/T2w ratio [26], the principal component of gene expression [13, 47], cerebral blood flow, and metabolic glucose uptake [72], whereas the other is composed of maps like the principal functional gradient [46], intersubject functional variability [51], and developmental and evolutionary expansion [34]. In both cases, these clusters recapitulate a number of associations previously reported in the literature [13, 26, 31, 67].

Functionality for easing the computation of and standardizing such comparisons in the future is necessary to ensure that new datasets can be integrated into our broader understanding of the human brain. The `neuromaps` toolbox offers just such functionality, including workflows to transform, compare, and contextualize brain maps. The `neuromaps` distribution contains high-quality, group-level transformations between four standard coordinate systems, uniform interfaces for comparing brain maps, and access to nine null models for use in generating statistical inferences. By developing the toolbox openly on GitHub (https://github.com/netneurolab/neuromaps), it is our hope that `neuromaps` can serve as a community tool for broad use in brain mapping research moving forward.

One consideration researchers must be aware of when using the `neuromaps` toolbox is that the provided transformations between coordinate systems are meant to be applied to group-level data; however, in general, when subject-level data are available it is better to reprocess them in the desired coordinate system rather than transforming group-level aggregate data. Unfortunately, in practice, subject-level data for many commonly-used brain maps are not available to researchers, and so having high-quality transformations between systems is critical to ensuring that analyses are performed in the most accurate manner possible. We have based the provided transformations on state-of-the-art frameworks (i.e., registration fusion and multimodal surface matching) which have been rigorously assessed and validated on other datasets [62, 63, 90]. Moving forward, as new frameworks arise for mapping between coordinate systems we will endeavor to provide updated transformations when possible.

Altogether, the current report introduces a new opensource Python package, `neuromaps`, for use in human brain mapping research. The toolbox offers researchers access to a wide repository of brain maps taken from the published literature, high-quality transformations between four standard coordinate systems, and uniform interfaces for statistical comparisons between brain maps. As the rate at which new brain maps are generated in the field continues to grow, we hope that `neuromaps` will provide researchers a set of standardized workflows for better understanding what these data can tell us about the human brain.

## ACKNOWLEDGEMENTS

This research was undertaken thanks in part to funding from the Canada First Research Excellence Fund, awarded to McGill University for the Healthy Brains for Healthy Lives initiative. This work was also supported in part by funding provided by Brain Canada, in partnership with Health Canada, for the Canadian Open Neuroscience Platform initiative. RDM acknowledges support from the Fonds du Recherche Québec - Nature et Technologies and the Canadian Open Neuroscience Platform. JYH acknowledges support from the Helmholtz International BigBrain Analytics & Learning Laboratory, the Natural Sciences and Engineering Research Council of Canada, and the Fonds du ResercheQuébec - Nature et Technologies. SB acknowledges support from the NIH (R01 EB026299), a Discovery Grant from the Natural Science and Engineering Research Council of Canada (436355-13), the CIHR Canada Research Chair in Neural Dynamics of Brain Systems. TDS acknowledges support from the NIH (R01 MH112847 and R01 MH120482). MMC acknowledges support from the Natural Sciences and Engineering Research Council of Canada, the Canada Research Chairs Program, Healthy Brains for Healthy Lives, and the Fonds du Recherche Québec - Nature et Technologies. BM acknowledges support from the Natural Sciences and Engineering Research Council of Canada (NSERC Discovery Grant RG-PIN #017-04265) and from the Canada Research Chairs Program.

---

**Table S1.** Standard coordinate system files available in neuromaps.

| *Coordinate system* | *Density / resolution* | *Provided files* |
| --- | --- | --- |
| fsLR | 4k | midthickness, inflated, sphere |
| fsLR | 8k | midthickness, inflated, sphere |
| fsLR | 32k | midthickness, inflated, very inflated, sphere |
| fsLR | 164k | midthickness, inflated, very inflated, sphere |
| fsaverage | 3k | white, pial, inflated, sphere |
| fsaverage | 10k | white, pial, inflated, sphere |
| fsaverage | 41k | white, pial, inflated, sphere |
| fsaverage | 164k | white, pial, inflated, sphere |
| CIVET | 41k | white, midthickness, inflated, very inflated, sphere |
| MNI152 | 1mm | T1w, T2w, PD, brainmask |
| MNI152 | 2mm | T1w, T2w, PD, brainmask |
| MNI152 | 3mm | T1w, T2w, PD, brainmask |

**Table S2.** Published brain maps available in neuromaps.

| *Map name* | *Coordinate system* | *Density* | *Citation* |
| --- | --- | --- | --- |
| Myelin map | fsLR | 32k | Glasser et al. [25] |
| Cortical thickness | fsLR | 32k | Glasser et al. [25] |
| Allen Human Brain Atlas PC1 | fsaverage | 10k | Markello et al. [47] |
| Delta power | fsLR | 4k | Van Essen et al. [78] |
| Theta power | fsLR | 4k | Van Essen et al. [78] |
| Alpha power | fsLR | 4k | Van Essen et al. [78] |
| Beta power | fsLR | 4k | Van Essen et al. [78] |
| Low gamma power | fsLR | 4k | Van Essen et al. [78] |
| High gamma power | fsLR | 4k | Van Essen et al. [78] |
| Intrinsic timescale | fsLR | 4k | Van Essen et al. [78] |
| Developmental expansion | fsLR | 164k | Hill et al. [34] |
| Evolutionary expansion | fsLR | 164k | Hill et al. [34] |
| Functional gradient | fsLR | 32k | Margulies et al. [46] |
| Inter-subject functional variability | fsaverage | 1k | Mueller et al. [51] |
| NeuroSynth PC1 | MNI152 | 2mm | Yarkoni et al. [92] |
| Cerebral blood flow (CBF) | fsLR | 164k | Vaishnavi et al. [72] |
| Cerebral blood volume (CBV) | fsLR | 164k | Vaishnavi et al. [72] |
| Oxygen metabolism (CMRO2) | fsLR | 164k | Vaishnavi et al. [72] |
| Glucose metabolism (GMRGlu) | fsLR | 164k | Vaishnavi et al. [72] |
| Allometric scaling (PNC) | CIVET | 41k | Reardon et al. [60] |
| Allometric scaling (NIH) | CIVET | 41k | Reardon et al. [60] |
| Neurotransmitter receptors/transporters | MNI152 | — | Hansen et al. [32] |

